# Epidemiology of Childhood Blindness: a Community based study in Bangladesh

**DOI:** 10.1101/532655

**Authors:** A.H.M Enayet Hussain, Junnatul Ferdoush, Saidur Rahman Mashreky, AKM Fazlur Rahman, Nahid Ferdausi, Koustuv Dalal

## Abstract

This study has aimed to detect the prevalence and causes of childhood blindness in rural area of Bangladesh. A cross sectional quantitative study design was adopted for this study which carried-out in three unions (sub-districts) of Raiganj upazila of Sirajganj district of Bangladesh. Using a validated tool a screening program was conducted at household level. After initial screening a team of ophthalmologists confirmed the diagnosis by clinical examinations. The prevalence of childhood blindness found was 6.3 per 10,000 children. The rate of uni-ocular blindness was 4.8 per 10,000 children. Congenital problems were the major cause of both uniocular and binocular blindness (UOB: 84% and BB: 92% &). For binocular blindness, whole globe was the responsible site (28.0%, CI: 13.1, 47.7) and cornea was for the uni-ocular blindness (57.8%, CI: 35.3, 78.1). Childhood blindness is a public health problem in Bangladesh. Childhood blindness is common irrespective of gender. Major causes of childhood blindness are congenital.

## Background

Globally, an estimated number of 36 million inhabitants are residing with blindness [1]. The prevalence by age distribution showed that around 1.4 million children aged 0-14 years are living with blindness while about 17.5 million are at risk of low vision [2]. An estimated 70 million blind-person years are caused by blindness among children [3]. Although the actual number of blind children is much lower than the blind adults, but the number of blind years resulting from the blindness is alarmingly high in children which has an immense social and economic impact [4–6].

The magnitude and causes of visual impairments and blindness varies from region to region because of socio-developmental diversification [4]. Analysis of global data showed that around 90% of the blind people belongs to the developing countries [7]. There is dearth of recent population based studies showing the prevalence as well as factors responsible for childhood blindness in the context of developing countries. However, data found that the burden of childhood blindness higher in African and Asian region which is mainly due to inaccessible primary health care services [5,8,9].

Majority of the causes for childhood blindness are avoidable with the available minimal resource settings of the developing countries [7]. Being a lower-middle income country, Bangladesh is not the exception. The situation is more miserable in the rural context. Rural areas in Bangladesh are already facing enormous healthcare delivery problem [10]. With socioeconomic and cultural constraints and with the backdrop of medical poverty-trap, it could be assumed that Bangladesh, especially rural areas have poor healthcare facilities for eye care services [11]. Over the last few years, number of initiatives have been taken to achieve the VISION-2020. [12,13]. However, there is further scope of improvement in order to achieve the goal. Moreover, there is scarcity of data from the existing researches related to the burden and capacity of health system regarding childhood blindness. This study aims to detect the prevalence and causes of childhood blindness at community level in rural area of Bangladesh.

## Methods

### Study Design

A cross sectional quantitative design was adopted for this study, carried-out during January and April 2017 in three unions of Raiganj, a sub-district (Upazila) of Sirajganj District in the Rajshahi Division of northern Bangladesh.

### Study Population

The whole area of the targeted upazila is 259.74 sq. km with an established population density of 223 per sq. km. About 317,666 people are inhabiting in the upazila where male and female proportion were 51.1% and 49.9% respectively [14]. The Upazila comprises of nine unions. A union is the lowest administrative unit comprising of 20,000-50,000 population. The survey was conducted among three unions where Center for Injury Prevention and Research Bangladesh (CIPRB) is maintaining an injury and demographic surveillance system. Entire population of those three unions are included in this surveillance system of the current study. It comprised 31,971 households with a population of 147,072 where total number of children aged ≤15 years was 39,351. The survey was conducted among whole child populations of the indicated study areas. The target population was below or of 15 years of old child included according to the operational definition.

### Data Collection Procedure

A household level screening was conducted to identify the suspected childhood blindness, visual impairment and other ocular morbidities. In the screening, data collectors conducted face-to-face interview along with some simple examinations using a validated structured questionnaire with pictorial charts. Each data collector was provided training on measuring visual acuity using age specific vision chart and performing of a basic ocular examination. A day long comprehensive training was provided by a team of both the researchers and ophthalmologists. Socio-demographic information of the respondents was collected from the database of CIPRB surveillance system. All individuals in the surveillance area have their own unique identification number. After collecting specific information, it was merged with the required socio-demographic variables of the existing population database.

Total 18 field workers (3 field supervisors and 15 data collectors) conducted the survey. If any household member was not available during the period of data collection, field workers collected their contact number and visited them again as per their convenience. Data collectors identified 570 suspected cases blindness, visual impairment and other ocular morbidities.

All the suspected children were then invited to visit the eye camp for the confirmation of their diseases and further management advice. Field workers invited all the parents/ caregivers of the suspected children through mobile phone, those who were not reached out over phone in-person meeting was carried out to ensure presence in the eye camp. Transportation cost were given to all families. Day-long eye camp was carried out in three phases to examine all suspected cases. Team was consisting of five ophthalmologists, two health assistants and five supporting staffs.

All children were examined as per the standard clinical guideline after having written consent from their legal guardian. Ophthalmologists used Snellen chart, slit lamp, retinoscope, direct ophthalmoscope and indirect ophthalmoscope to confirm the cases. Out of 570 screened cases finally, 198 cases of blindness and different kind of eye morbidities were confirmed by the team. The ophthalmologist kept the record of both the principal reason as well as the contributing reasons for their blindness and visual impairment. Among the confirmed cases, total nine children were referred to the specialized hospitals for carrying out their indicative surgery. Other confirmed cases were referred to different health care facilities for definitive treatment (Fig 1).

**Fig 1:**
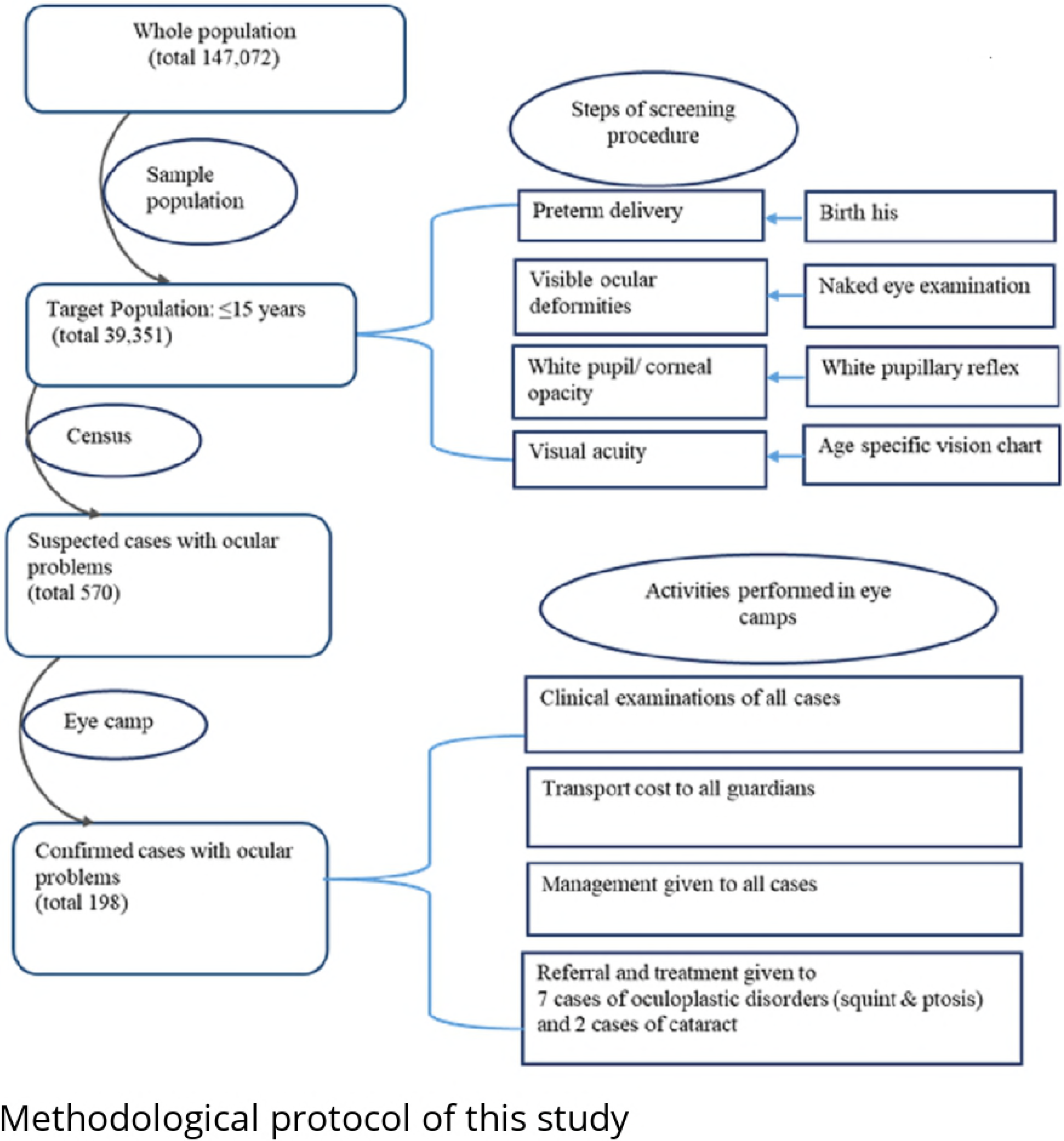
Methodological protocol of this study

### Statistical Analysis

Descriptive analysis was carried out to describe the population by gender, age and socioeconomic status. Similar analysis was made to estimate the prevalence of childhood blindness and its associated risk factors. The prevalence rate of all types of blindness were calculated per 10,000 children with 95% confidence interval (CI). Prevalence of childhood blindness was described by gender, age, wealth index and classification of the blindness. WHO standard anatomical and aetiological classification used to do the analysis [15]. For analysis purpose, child age group was categorized as under-five and 5 to 15 years where wealth index categorized as poor, middle and rich. The wealth index was categorized using some selective assets of the household which included unit of land ownership; availability of amenities i.e. electricity, refrigerator, television, radio, bicycle, motorcycle, wardrobe, table, chair, clock, bed, sewing machine, mobile, car, water, toilet; daily used fuels; household income along with expenditure and the materials used for building roof, floor and wall of the household. Bi-variate analysis was carried out to analysis the relationship between childhood blindness and the independent variables such as sex, age group, wealth index and classification of blindness. Construction of all variables and estimations were ascertained using the statistical software SPSS version 24.

### Operational Definitions

#### Children

In this study, cases at or below 15 years of age were considered as children.

#### Blindness

Blindness definition of ICD-10 considered for this study. Blindness is the corrected visual acuity of less than 3/60 in the better eye or a central field of less than 10 degrees [16].

#### Binocular and Uniocular Blindness (BB & UOB)

ICD-10 standard definition of Binocular and Mono/Uni ocular blindness considered for this study [16].

#### Reversible Blindness

The temporary loss of eye sight which could be treatable through surgical intervention is termed as reversible blindness.

#### Irreversible Blindness

Irreversible blindness where the eye sight can be restored either medical or surgical intervention.

### Ethical Consideration

Ethical clearance of this study was received from the Ethical Review Committee of Centre for Injury Prevention and Research Bangladesh. We have taken written consent of all legal guardians of the respective suspected cases during the screening and for the ophthalmologist visit.

## Results

In this study, the proportion of male (51%) and female (49%) population was almost similar. Two categories of age group for the study population were generated and majority of the children were from age group 5 to 15 years (70%) and rest were below 5 years old. Four categories of wealth index were generated for the study population and majority of the children have fallen into the poor category (50%) (Table 1).

**Table 1:** Population characteristics (N= 39,351)

| Variables | Frequency | Percentage |
| --- | --- | --- |
| **Gender** |  |  |
| Male | 20211 | 51.3 |
| Female | 19140 | 48.6 |
| **Age (in years)** |  |  |
| < 5 years | 11806 | 30.0 |
| 5 to 15 years | 27545 | 70.0 |
| **Wealth index** |  |  |
| Poor | 19678 | 50.0 |
| Middle-class | 9843 | 25.0 |
| Rich | 9830 | 25.0 |

The prevalence of binocular blindness among study population found was 6.3 per 10,000 children whereas the rate for uni-ocular blindness found was 4.8 per 10, 000 children. Both uni-ocular and binocular rate was found higher among male, however difference was not statistically significant. Significantly higher rate of binocular was found among the under 5 aged children. The rates were 19.4 (CI: 12.6 - 28.7) and 0.7 (CI: 0.1 - 2.3) in under-five and 5 years and above age group children. Compared to rich higher rate of binocular blind was found among poor.

However, difference was not statistically significant. The rates were 8.1 (CI: 4.7 - 12.9) and 2.0 (CI: 0.3 - 6.7) among rich and poor respectively (Table 2).

**Table 2:**
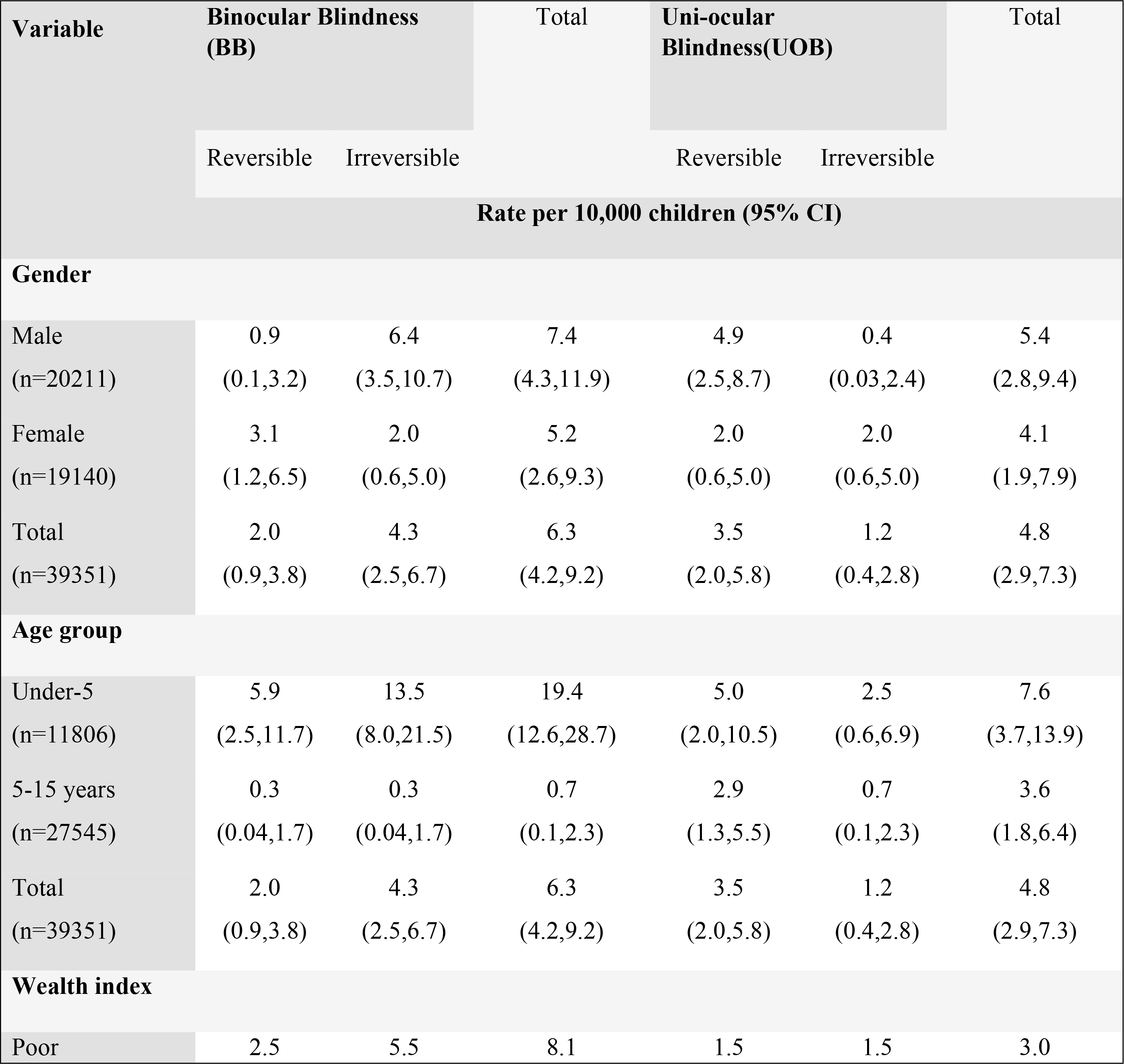
Distribution of BB & UOB by Age, Sex and Socio-economic Status

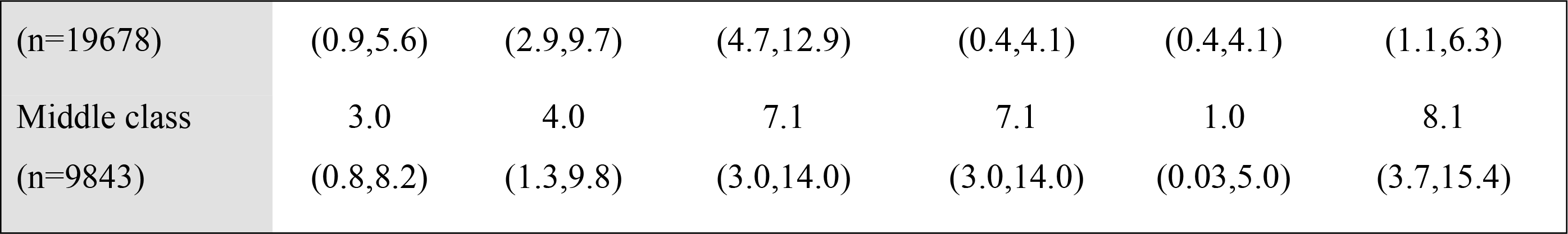

Congenital diseases are found as the major reason for both the uniocular and binocular blindness (BB: 92% & UOB: 84%). For binocular blindness, whole globe was the responsible site for structural origination with the proportion of 28.0 (CI: 13.1 - 47.7) where cornea was for uni-ocular blindness (57.8%, CI: 35.3 - 78.1) (Table 3).

**Table 3:**
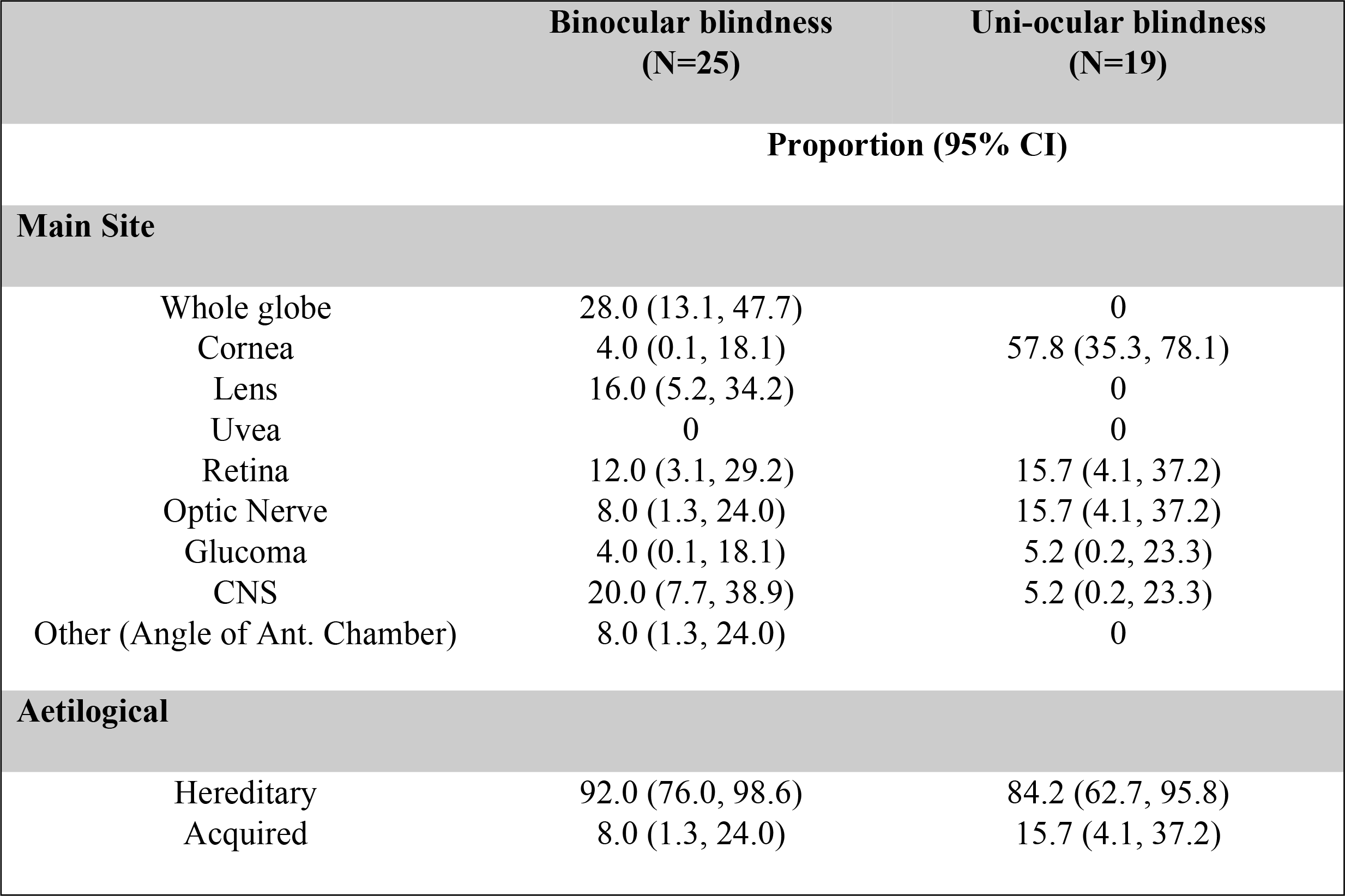
Distribution of BB & UOB by anatomical & aetiological classification

## Discussion

This is the first household level survey in a rural community of Bangladesh showing the prevalence of childhood blindness. The rate was 6.3 per 10,000. By extrapolating this number into the national target group, it is found that around 35,000 children in Bangladesh is living with blindness. While another extrapolation of the data found from a study of Muhit et al. in 2007 claimed that the estimated children with blindness/ severe visual impairments could be 26,000 [17]. Both studies followed a different methodology where the former screened out the cases through house to house community based survey and the latter followed a combination of the inclusion of special child from the institution along with capturing of blind cases through key informant method. Moreover, another assumption made that around 40,000 children of Bangladesh are breathing their lives with blindness [18]. It is mentionable that the information of the current study was extracted from the rural context only, warranting further rigorous research in both rural and urban context.

This is the very first study which estimated the prevalence and causes of uni-ocular reversible and irreversible blindness among children. There was almost no previous research found in terms of Bangladesh that examined the burden and etiology of uni-ocular blindness. Data from a community based study of Oman showed that the rate of uni-ocular blindness is 9 per 10,000 children [19]. The estimation of blindness will be double if the contributory number of uni-ocular blindness dropped into the bucket of binocular blindness [19]. So, this piece of information might be helpful for the policy makers to set their priorities. Study stated that the persistent untreated unilateral blindness particularly due to ocular trauma eventually leads to blindness [20]. Although the prevalence of uni-ocular blindness does not directly contribute to the burden of total blindness, but early diagnosis and treatment of reversible uni-ocular blindness will eventually contribute in reducing the burden of total blindness. By that means, the burden of uni-ocular blindness is also an emerging public health concern for Bangladesh.

Study stated majority of blind children are female as found in t population-based study at two areas of Andhra Pradesh, India which was published in 2003 [21] In our study, we found higher rate among male however it is not statistically significant.. The relative small sample size should keep in consideration while doing the interpretation of this study finding. Though many studies showed that blindness is highest among poor group than the richest but this study does not resemble such pattern [22]. Study finding of this study showed the similar trend although the difference was not statistically significance.

Congenital causes were found as the most common reason for both uni-ocular and binocular childhood blindness. Additionally, whole globe was the responsible site in terms of binocular blindness whereas cornea found as the accountable for mono ocular blindness. A previous nationwide study stated that lens as the commonest anatomical site where unoperated cataract cases were the responsible reason [23]. Improved maternal and child care initiatives taken by the Bangladesh Govt. might help in reducing national burden of cataract in children. A study conducted in school going children of China also found whole globe as the accountable anatomical site [24]. Previous study result showed Vitamin A deficiency as the major attributable acquired reason for childhood vision loss [23]. Nevertheless, this study did not find these acquired reasons to be as mentionable which might be due to continuous spectacular progress in the sector of health and nutrition in Bangladesh. The National Vitamin-A plus campaign is a successful program held each year in Bangladesh which contributed tremendously to control night blindness [25].

### Strength and limitation

This study has performed screening of the cases at community level through house to house survey following another study published in 2003 done by Nirmalan et al in Southern India [26]. Investigators suggest that community door to door approach is credible to determine the extent of childhood blindness as there is less chance of case dropping. Moreover, this survey was conducted within an established *Health and Demographic Surveillance System* which covered more than 150,000 population. All the children aged below and 15 years of the study area were included in this study. It minimized the sampling error of the study. A group of senior ophthalmologists were involved in the diagnostic procedure. An experienced ophthalmologist leaded the diagnostic team. It might minimize the error related to the misclassification of diseases. The study was conducted in a rural area of Bangladesh, which may not represent the urban situation of the country.

### Conclusion

Childhood blindness is a significant public health concern in Bangladesh. Childhood blindness is common irrespective of gender. Major causes of childhood blindness are congenital.

### Recommendation

This study finding indicates that necessary measures are needed to be considered for the reduction of childhood blindness in Bangladesh. Raising social awareness is important so that the reversible childhood blindness can be identified and treated early. A strategy need to be developed with proper strengthening of six building blocks of the health system for the prevention of childhood blindness in Bangladesh.

## Acknowledgements

This study was conducted in the surveillance area of Centre for Injury Prevention and Research Bangladesh (CIPRB). CIPRB provided administrative and technical support in the data collection and data management procedure. Disabled Rehabilitation Research Association (DRRA), Bangladesh provided financial support for this study. We are also grateful to the ophthalmologists who took part in the visits to detect the confirmed cases. We are thankful to Director General of Health Services (DGHS) for providing their technical support in validating the materials.

## Authors Contributions

Conceptualization: EH, SRM, KD

Formal Analysis: EH, JF, SRM, KD

Writing (original draft): EH, JF, SRM, NF, KD

Writing (review and editing): EH, SRM, FR, KD

